# Structure of the human heterotetrameric *cis*-prenyltransferase complex

**DOI:** 10.1101/2020.06.02.095570

**Authors:** Michal Lisnyansky Bar-El, Pavla Vankova, Petr Man, Yoni Haitin, Moshe Giladi

## Abstract

The human *cis*-prenyltransferase (h*cis*-PT) is an enzymatic complex essential for protein N-glycosylation. Synthesizing the precursor of the glycosyl carrier dolichol-phosphate, we reveal here that h*cis*-PT exhibits a novel heterotetrameric assembly in solution, composed of two catalytic dehydrodolichyl diphosphate synthase (DHDDS) and two inactive Nogo-B receptor (NgBR) subunits. The 2.3 Å crystal structure of the complex exposes a dimer-of-heterodimers arrangement, with DHDDS C-termini serving as homotypic assembly domains. Furthermore, the structure elucidates the molecular details associated with substrate binding, catalysis, and product length determination. Importantly, the distal C-terminus of NgBR transverses across the heterodimeric interface, directly participating in substrate binding and underlying the allosteric communication between the subunits. Finally, mapping disease-associated h*cis*-PT mutations involved in blindness, neurological and glycosylation disorders onto the structure reveals their clustering around the active site. Together, our structure of the h*cis*-PT complex unveils the dolichol synthesis mechanism and its perturbation in disease.

## Introduction

Prenyltransferases are essential enzymes that synthesize isoprenoids, an enormous group of chemically diverse compounds participating in a myriad of cellular processes in all living cells (Grabińska et al., 2016). With chain lengths varying from C_10_ (geranyl diphosphate) to >C_10,000_ (natural rubber), isoprenoids are synthesized by chain elongation of an allylic diphosphate primer via a variable number of condensation reactions with isopentenyl pyrophosphate (IPP, C_5_) (Ogura and Koyama, 2002; Takahashi and Koyama, 2006). Prenyltransferases are classified as *cis*-prenyltransferase or *trans*-prenyltransferase according to the double bonds they form during the condensation reaction (Ogura and Koyama, 2002). *Cis*-prenyltransferases are further classified according to their product chain length into short-chain (C_15_), medium-chain (C_50-55_), long-chain (C_70-120_), and rubber synthases (Grabińska et al., 2016). Importantly, while short- and medium-chain *cis*-prenyltransferase complexes are homodimeric, long-chain *cis*-prenyltransferases and rubber synthases are formed by a heteromeric subunit assembly of unknown stoichiometry (Grabińska et al., 2016; Yamashita et al., 2016). To date, only homodimeric enzymes were structurally characterized (Fujihashi et al., 2001; Guo et al., 2005; Ko et al., 2019; Wang et al., 2008). Therefore, our understanding of the mechanisms allowing long-chain isoprenoid formation by heteromeric enzymes remains limited.

The human *cis*-prenyltransferase complex (h*cis*-PT) catalyzes the formation of dehydrodolichyl diphosphate (DHDD, C_85-100_), a long-chain isoprenoid, by chain elongation of farnesyl diphosphate (FPP, C_15_) via multiple condensations with IPP (Figure 1A) (Harrison et al., 2011). DHDD is the precursor for dolichol-phosphate, the lipidic glycosyl carrier crucial for N-linked protein glycosylation (Figure 1A) (Schwarz and Aebi, 2011). Localized to the endoplasmic reticulum, h*cis*-PT is composed of two structurally and functionally distinct subunit types. These include the catalytically-active DHDD synthase (DHDDS) and the quiescent Nogo-B receptor (NgBR) subunits (Harrison et al., 2011). Importantly, while DHDDS subunits are cytosolic, NgBR can be subdivided into an N-terminal transmembrane domain and a C-terminal pseudo *cis*-prenyltransferase domain, which lacks detectable catalytic activity and directly interacts with DHDDS (Figure S1) (Harrison et al., 2011). We have previously shown that DHDDS can form functional homodimers, but these complexes exhibit poor catalytic activity compared to the homodimeric orthologs or the heteromeric h*cis*-PT (Grabińska et al., 2017; Guo et al., 2005; Lisnyansky Bar-El et al., 2019). Accordingly, previous studies suggested that NgBR can allosterically modulate the activity of the catalytic DHDDS subunit (Grabińska et al., 2017; Harrison et al., 2011; Park et al., 2014). Indeed, overexpression of NgBR was shown to significantly enhance h*cis*-PT activity in cells, supporting an NgBR-mediated allosteric modulation of DHDDS activity (Harrison et al., 2011; Park et al., 2014). Although the quest for elucidating the functional and structural roles of NgBR activity in the context of the h*cis*-PT complex is still ongoing, this effect was suggested to involve a conserved RxG motif, localized to the NgBR C-terminal tail (Grabińska et al., 2017), but the underlying mechanism remains to be determined.

**Figure 1.**
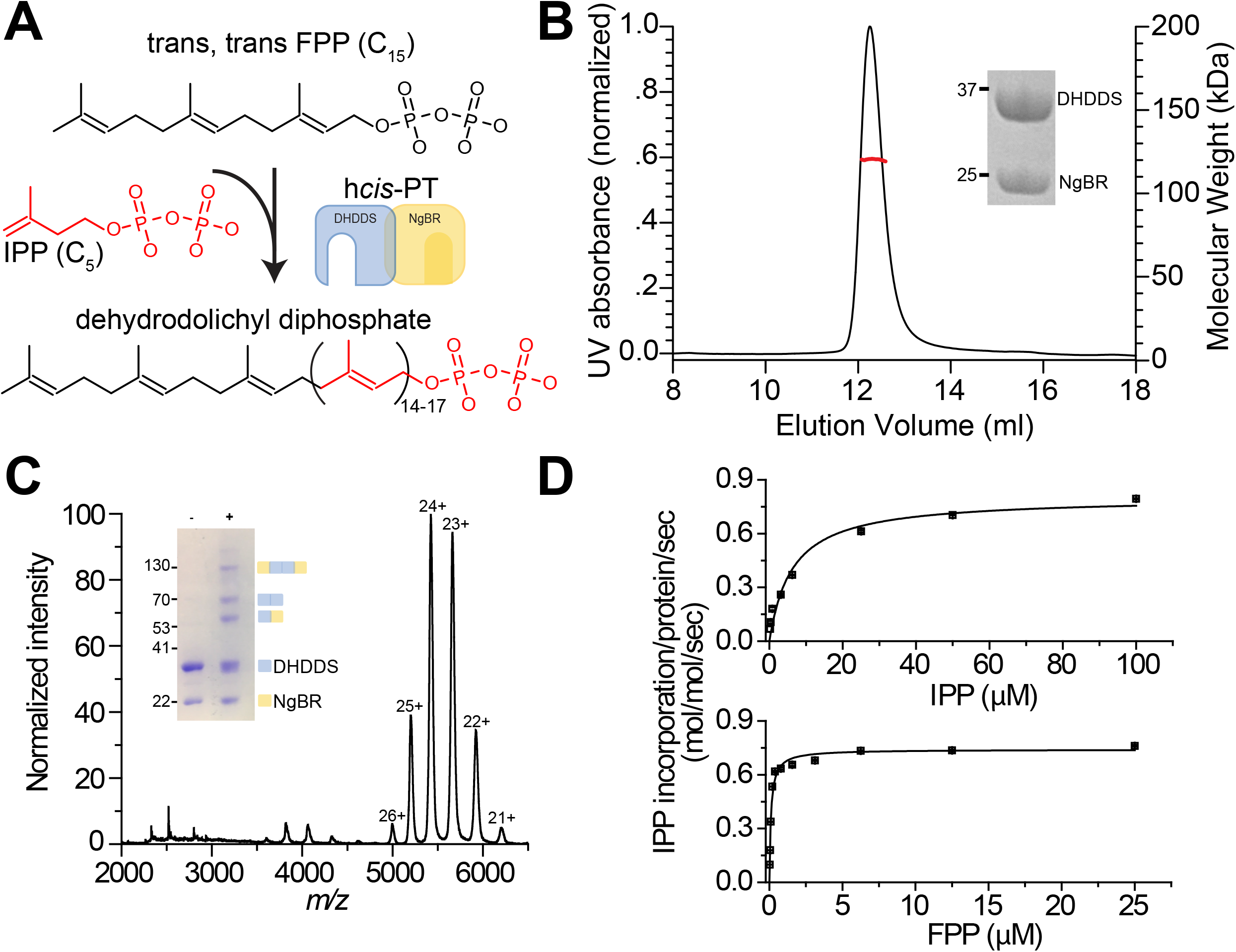
Complex stoichiometry and functional analysis of sh*cis*-PT. **(A)** Dehydrodolichyl diphosphate synthesis reaction scheme. **(B)** SEC-MALS analysis of the purified sh*cis*-PT. The black and red curves indicate UV absorption and molecular weight, respectively. Inset: SDS-PAGE analysis of the purified complex. **(C)** Native ESI-MS spectrum obtained using low activation conditions. The main distribution corresponds to the heterotetramer (charge states 21-26). Inset: SDS-PAGE analysis of the complex following glutaraldehyde cross-linking. **(D)** *In vitro* activity of purified sh*cis*-PT, assessed as IPP incorporation. Experiments were performed as described in the methods section. Data are presented as mean ± SEM (n = 3).

In line with the crucial significance of N-linked glycosylation for proper cellular function, mutations in both h*cis*-PT subunits were associated with human diseases. Specifically, DHDDS missense mutations were shown to result in phenotypes ranging from autosomal recessive retinitis pigmentosa (Zelinger et al., 2011; Züchner et al., 2011), through developmental epileptic encephalopathies (Hamdan et al., 2017), to a case of fatal congenital disorder of glycosylation reported in a patient heterozygous for both a splice site mutation and a nonsense mutation (Sabry et al., 2016). Moreover, a missense mutation in the conserved C-terminal RxG motif of NgBR was shown to cause a congenital glycosylation disorder with refractory epilepsy, visual and neurological impairments, congenital scoliosis, and hearing deficit (Park et al., 2014). Finally, recently identified missense mutations in NgBR were shown to contribute to the etiology of Parkinson’s disease (Guo et al., 2018).

Despite an immense body of work focusing on the biochemical and structural properties of *cis*-prenyltransferases, our mechanistic understanding of these enzymes arises mainly from investigations of homodimeric prokaryotic orthologs (Takahashi and Koyama, 2006). Thus, in contrast with the growing clinical relevance of h*cis*-PT, the basic mechanisms underlying its heteromeric assembly, intersubunit communication and long-chain isoprenoid synthesis remain poorly understood. Here, we established an overexpression and co-purification paradigm of the h*cis*-PT complex devoid of its transmembrane region. Unexpectedly, the purified complex formed stable heterotetramers in solution, with equimolar stoichiometry of DHDDS and NgBR subunits. Moreover, it exhibited a marked activity enhancement compared with the purified homodimeric DHDDS (Lisnyansky Bar-El et al., 2019). Next, we determined the co-crystal structure of the complex with bound FPP, phosphate and Mg^2+^ at 2.3 Å resolution. Strikingly, while DHDDS encompasses an active site reminiscent to that observed in homodimeric family members, the structure exposes the long-sought involvement of NgBR in active site organization and provides insights into the molecular mechanisms associated with the functional enhancement it confers. Moreover, it reveals the structural organization of the DHDDS C-terminal domain, exhibiting a ‘helix-turn-helix’ motif fold never observed before in other *cis*-prenyltransferases, which facilitates complex tetramerization via a dimer-of-heterodimers assembly mode. Finally, the structure lays the foundation towards identifying the determinants governing long-chain isoprenoid synthesis.

## Results

### Purification and *in vitro* activity characterization of the soluble h*cis*-PT complex

Previously, sequence and biochemical analyses of NgBR revealed that it interacts with DHDDS via its cytosolic C-terminal pseudo *cis*-prenyltransferase homology domain (Figure S1) (Harrison et al., 2011). Thus, we generated an NgBR construct solely encompassing its cytosolic domain (sNgBR, residues 73*-293*, where the asterisks designate NgBR residues). Importantly, previous studies using yeast complementation showed that truncation up to position 85* did not affect the ability of NgBR to support cell growth following co-transformation with DHDDS, indicating the catalytic function of the complex is preserved in the absence of the transmembrane domain (Grabińska et al., 2017). Next, we co-overexpressed the full-length human DHDDS (residues 1-333) and sNgBR in *E. coli*. Following purification, we obtained a homogeneous population of heteromeric soluble h*cis*-PT (sh*cis*-PT) (Figure 1B).

Intriguingly, during the final size-exclusion purification step, we noticed that the elution volume of sh*cis*-PT corresponds to a higher than expected molecular weight range. Strikingly, size-exclusion chromatography multi-angle light-scattering (SEC-MALS) analysis of the purified sh*cis*-PT revealed a monodispersed population with a molecular weight of 119.7 ± 0.4 kDa (Figure 1B). Within the experimental error of SEC-MALS, this mass may correspond to a stable heterotetramer composed of either two DHDDS (monomer molecular weight = 39.1 kDa) and two sNgBR (monomer molecular weight = 25.2 kDa) subunits or one DHDDS and three NgBR subunits. In order to determine the stoichiometry of the complex, we used native electrospray ionization (ESI) mass-spectrometry (MS) (Figure 1C). Importantly, native ESI spectra, obtained at low activation conditions to preserve the tertiary structure, revealed a mass of 130.1 ± 0.2 kDa, confirming the heterotetrameric organization of the complex with a stoichiometry of two DHDDS and two sNgBR subunits (Figure 1C). However, under these conditions the deconvoluted mass was higher than predicted due to presence of sodium cations and low molecular weight adducts. Indeed, along with partial complex disintegration upon stepwise activation, a shift of the heterotetrameric population toward lower mass (128.7 ± 0.02 kDa) was observed and an agreement between theoretical and calculated masses was achieved (Figure S2). Finally, we used a cross-linking approach which enables the identification of subunits interaction within the heterotetramer. Treatment with glutaraldehyde, a homobifunctional amine-reactive crosslinker, resulted in the emergence of high-order oligomers, culminating in a heterotetrameric complex with a mass of ~125 kDa (Figure 1C). Additionally, although the cross-linking treatment can theoretically yield three types of dimers (DHDDS homodimer, sNgBR homodimer and DHDDS-sNgBR heterodimer), we could clearly detect only two bands, corresponding to DHDDS homodimers and DHDDS-sNgBR heterodimers, suggesting that the NgBR subunits exhibit spatial separation in the context of sh*cis*-PT. Together, the biophysical and biochemical characterizations of sh*cis*-PT suggests a dimer-of-heterodimers arrangement.

Next, in order to validate that the purified complex is catalytically viable, we tested its activity *in vitro* using a radioligand-based assay (Figure 1D). The purified sh*cis*-PT exhibited *k*_cat_ = 0.74 ± 0.02 s^−1^ and K_m_ = 6.23 ± 1.50 and 0.11 ± 0.01 μM for IPP and FPP, respectively. The *k*_cat_ value is similar to that previously reported for the intact h*cis*-PT (Grabińska et al., 2017) and ~400 fold higher compared to that of homodimeric DHDDS (Lisnyansky Bar-El et al., 2019). These results demonstrate that sh*cis*-PT recapitulates the function of the intact complex, further reinforcing that the absence of the N-terminus of NgBR does not impair the catalytic activity of the complex. Moreover, the increased activity of the heteromeric complex compared with homodimeric DHDDS points towards an allosteric mechanism of inter-subunit communication, since NgBR lacks catalytic activity (Grabińska et al., 2016).

### Structure overview of the sh*cis*-PT subunits

In order to unveil the structural basis of the functional allostery observed in the context of the sh*cis*-PT, we sought to determine its structure using X-ray crystallography. However, as initial crystallization attempts were unsuccessful, we removed residues 167-175 from NgBR (sNgBRΔ167*-175*), corresponding to an unresolved loop in the structure of the yeast homolog Nus1 (Ma et al., 2019), assuming that this region is highly flexible and precludes crystallization. Indeed, this construct (termed hereafter x*cis*-PT) resulted in well diffracting crystals, allowing us to determine its structure in complex with Mg^2+^ and FPP at 2.3 Å resolution. The asymmetric unit (ASU) contains a single heterodimer (Figure 2A), with Mg^2+^-FPP and an additional phosphate moiety bound only at the DHDDS active site.

**Figure 2.**
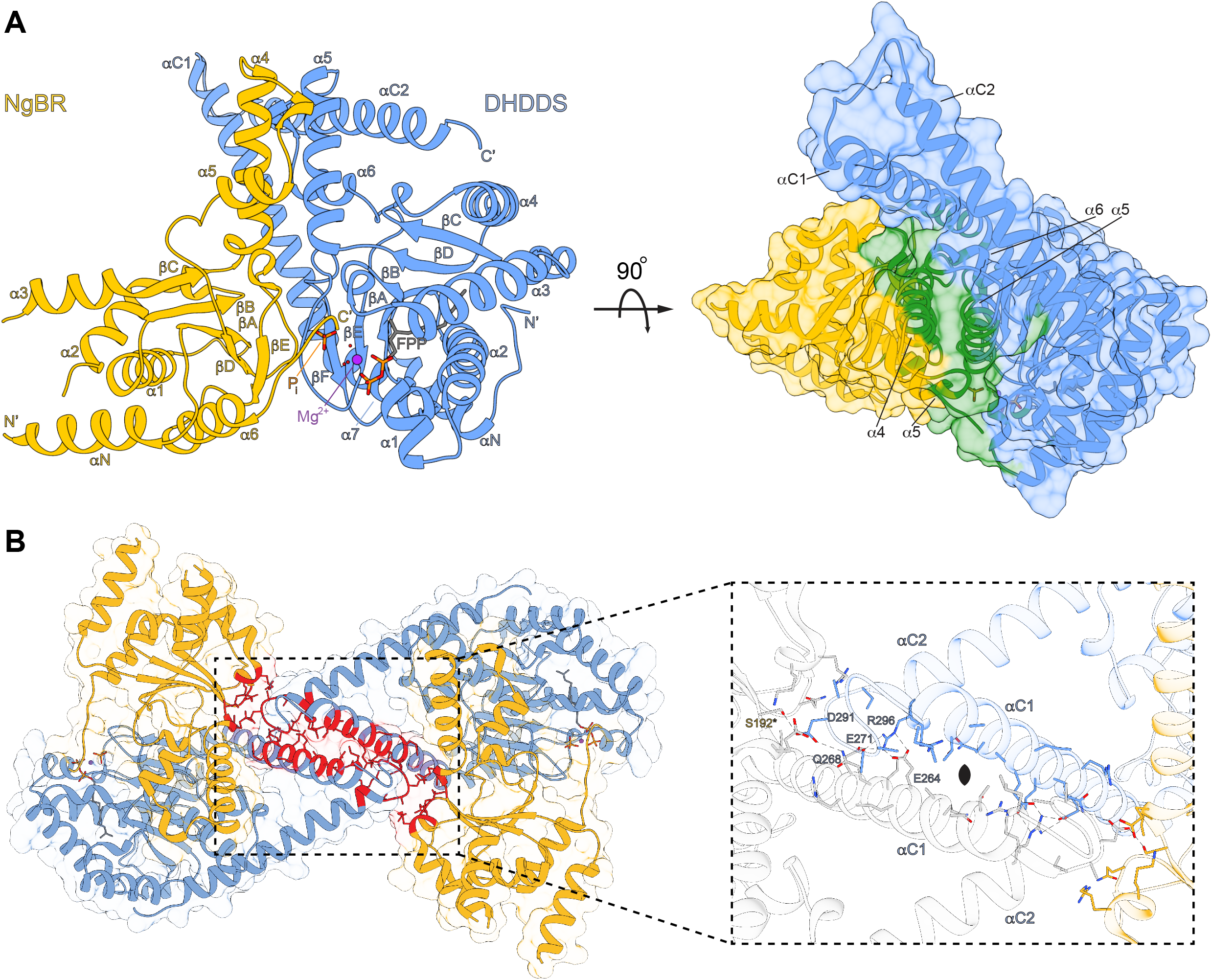
Overall structure of x*cis*-PT. **(A)** Left: cartoon representation the ASU, composed of a DHDDS-sNgBRΔ167-175 heterodimer. Secondary structure elements are labeled. Right: Surface representation of the ASU with residues involved in the heterodimeric interface highlighted in green. **(B)** The heterotetrameric complex assembly mode. Residues involved in tetramerization are shown as sticks and colored red. The rectangle frames a zoom perspective of the interactions between one heterodimer (colored as in panel A) and a heterodimer from the neighboring ASU (colored white).

DHDDS can be subdivided into three domains: (*i*) An N-terminal domain (NTD, residues 1-26); (*ii*) A canonical catalytic *cis*-prenyltransferase homology domain (residues 27-250); and, (*iii*) A C-terminal domain (CTD, residues 251-333) (Figure S1). The catalytic *cis*-prenyltransferase homology domain, which forms the central region of DHDDS and serves for heterodimerization with NgBR (Figure 2A, right), is composed of 7 α-helices and 6 β-strands (Figure S1), engulfing an elongated active site cavity. This domain shares high homology with undecaprenyl diphosphate synthase (UPPS), a bacterial medium-chain *cis*-prenyltransferase homolog, with r.m.s.d. = 0.76Å (Guo et al., 2005). The NTD and CTD flank the catalytic domain. The NTD is wrapped around the catalytic domain (Figure 2A, left), with its single α-helix, αN, directly packed against helix α7 via a network of hydrophobic interactions. Lastly, the CTD, absent from short- or medium-chain *cis*-prenyltransferases, features a novel ‘helix-turn-helix’ motif composed of two consecutive helices situated immediately downstream of α7 (Figures 2A, S1). The first helix, αC1, is kinked by 30 degrees relative to the preceding α7 from the catalytic domain. The second helix, αC2, is stabilized against αC1 by a salt-bridge between D273 and R306, and also a slew of electrostatic (E168-K320, E168-R321, D182-R309, R196-E318) and hydrophobic interactions with the heterodimerization interface between DHDDS and NgBR (Figure 2A).

Similar to DHDDS, the remnant N-terminal domain of sNgBR (NTD, residues 79*-100*) also encompasses a single α-helix, αN, likely serving as a structural link between the transmembrane and cytosolic domains in the intact protein. However, while NgBR was previously thought to share the canonical *cis*-prenyltransferase fold, similar to DHDDS (Harrison et al., 2011), the structure reveals that the two subunits share low structural similarity (r.m.s.d. = 2.07Å, residues 100*-293* of sNgBR and 27-250 of DHDDS). Indeed, the pseudo *cis*-prenyltranseferase homology domain of sNgBR encompasses only 6 α-helices and 5 β-strands (Figure 2A, S1), in contrast to the 7 α-helices and 6 β-strands found in the other *cis*-prenyltransferases (Takahashi and Koyama, 2006). This altered fold prevents interactions with substrates and hinders the catalytic activity of NgBR, as specified below.

### Mechanism of tetramerization via a dimer-of-heterodimers assembly

The DHDDS-sNgBR heterodimer is formed through a large interaction interface, with a buried surface area of 1938.0 Å^2^ (Figure 2A, right). This interface is mainly formed by helices α5, α6 and the βE-βF linker of DHDDS and helices α4, α5 and the βD-βE linker of NgBR, with an architecture reminiscent to that observed in homodimeric *cis*-prenyltransferases (Takahashi and Koyama, 2006). However, the heterodimeric interface features novel contacts between the NgBR C-terminus and the active site of DHDDS. Specifically, the structure reveals that the NgBR C-terminus, which encompasses the RxG motif and was previously suggested to play a critical role in h*cis*-PT activity, extends across the dimerization interface (Figure 2A, left). This transverse interaction results in its direct involvement in the organization of the active site of DHDDS and provides a structural framework for the intersubunit allosteric communication.

While crystal packing analysis suggested several possible biological assemblies, ranging from heterodimers to dodecamers, only one tetrameric assembly consistent with the oligomeric state in solution was detected. Importantly, this tetramer is formed by both homotypic interactions between DHDDS ‘helix-turn-helix’ motifs and heterotypic interactions of the ‘turn’ region with sNgBR from adjacent ASUs, burying a total surface area of 793.3 Å^2^ (Figure 2B). Specifically, this interface is stabilized by two polar networks: (*i*) A salt-bridges network, centered at the interaction between R296 from the αC2 helix of one heterodimer and E264 and E271 from αC1 helix of the adjacent heterodimer; (*ii*) A hydrogen-bonds network, originating from the interaction of D291, localized to the ‘turn’ between αC1 and αC2, and Q268 from αC1, with S192* of the adjacent heterodimer. Notably, this dimer-of-heterodimers arrangement is consistent with our cross-linking analysis (Figure 1C).

### Direct interaction between the RxG motif and catalytic residues in DHDDS

Although x*cis*-PT was crystallized in the presence of FPP and Mg^2+^, these substrates, along with an additional phosphate molecule, could only be detected in the active site of DHDDS (Figures 2A, 3). This active site is formed by a superficial polar region, stabilizing the interaction with the pyrophosphate headgroups, and a deep hydrophobic tunnel accommodating the elongating carbon chain (Figure 3A). The active site contains two substrate binding sites: An S_1_ site, which binds the initiatory substrate FPP, and an S_2_ site, which binds the IPP molecules used for chain elongation and is occupied by the phosphate molecule in our structure.

**Figure 3.**
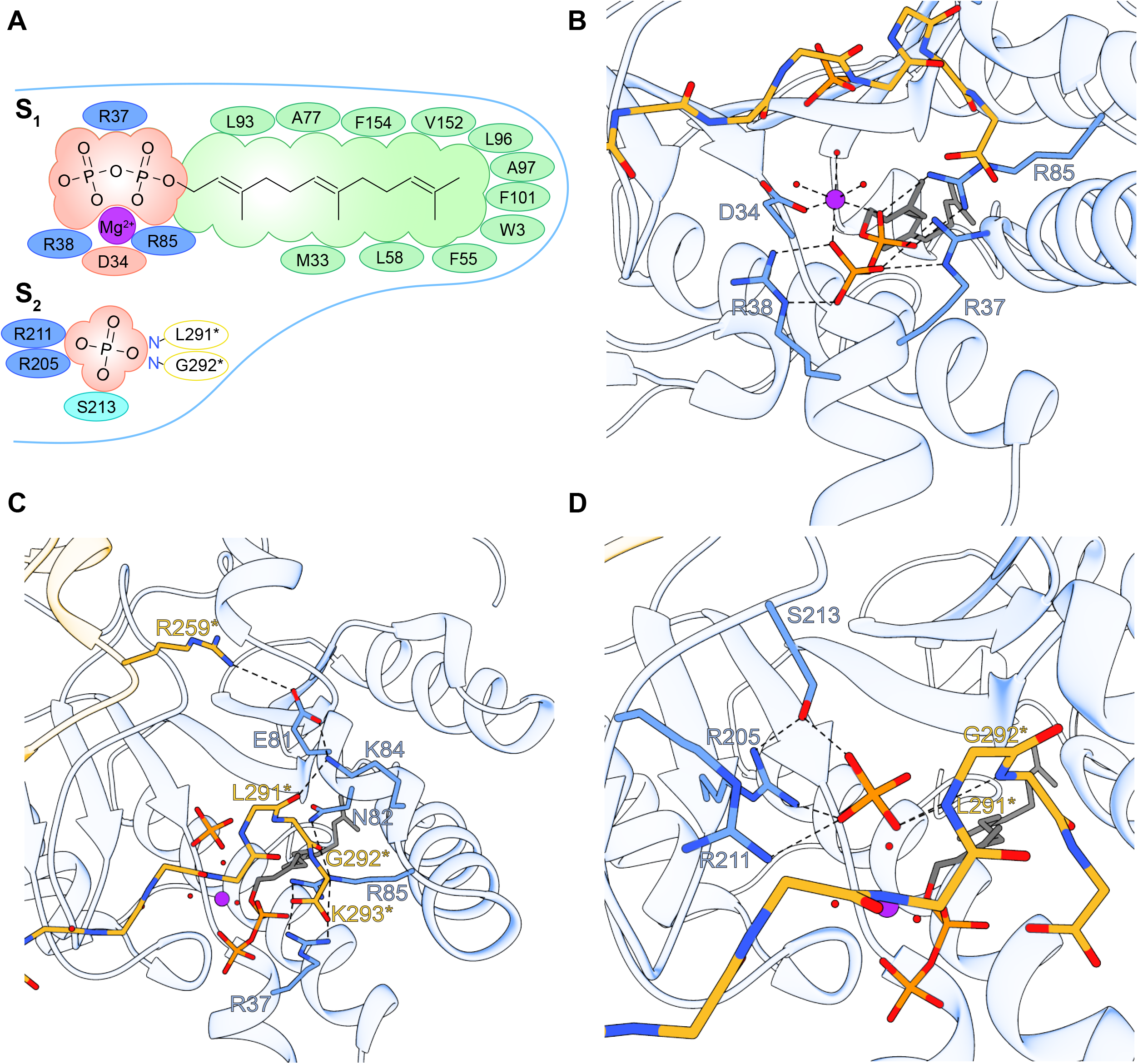
Structural organization of the active site. **(A)** Two-dimensional interactions diagram of the S_1_ and S_2_ sites. Hydrophobic interactions, positively and negatively charged DHDDS residues are colored green, blue and red, respectively. NgBR residues interacting with the S_2_ site via their backbone nitrogen are indicated in yellow circles. **(B)** The FPP pyrophosphate binding region at the S_1_ site. **(C)** Interactions of the NgBR C-terminal tail with the pyrophosphate binding region of the S_1_ site. **(D)** Phosphate coordination at the S_2_ site. Mg^2+^ and water molecules are shown as purple and red spheres, respectively. FPP and phosphate are shown as sticks.

At the S_1_ site, the pyrophosphate moiety of FPP interacts with a Mg^2+^ ion (Figure 3B). This Mg^2+^ ion, which plays a crucial role in pyrophosphate hydrolysis during the condensation reaction, is octahedrally coordinated by two of the pyrophosphate oxygens, three surrounding water molecules and one carboxylate oxygen of the strictly conserved D34 (Figure 3B) (Takahashi and Koyama, 2006). In addition to the interaction with the Mg^2+^ ion, the pyrophosphate moiety is also directly stabilized by R37 and R38 from helix α1, and R85 from the βB-α3 linker (Figure 3B).

Strikingly, the C-terminus of NgBR, encompassing the highly conserved and functionally important RxG motif (Grabińska et al., 2017), is directly involved in the spatial organization of the S_1_ site (Figure 3C). Specifically, it stabilizes the βB-α3 linker of DHDDS by forming two polar interactions networks, interweaved between NgBR and DHDDS. One network is organized such that two salt bridge interactions, between E81 and R259*, and K84 and the carbonyl oxygen of L291*, are anchored together by another salt bridge interaction between K84 and E81. The other network is formed by G292* backbone carbonyl interaction with N82 and R85 (Figure 3C), the latter involved in coordination of FPP (Figure 3B). In addition, K293*, forming the C-terminal carboxylate group of NgBR, interacts with both R37 and R85. Thus, our structure provides a plausible explanation for the high conservation of the RxG motif due to its role in active site organization.

While x*cis*-PT was crystallized in the absence of IPP, a phosphate molecule is occupying the IPP pyrophosphate group position, as observed in other *cis*-prenyltransferases (Figure S3), is present at the S_2_ site and is stabilized through a concerted coordination by the conserved R205, R211 and S213 (Figure 3D) (Takahashi and Koyama, 2006). In addition to the interactions of the NgBR C-terminus with the S_1_ site of DHDDS, the backbone nitrogen atoms of L291* and G292* also directly coordinate the phosphate molecule at S_2_ (Figure 3D). Together, the binding mode of the phosphate group suggests that h*cis*-PT and other *cis*-prenyltransferases probably share a similar S_2_ site.

### Hydrophobic interactions in the active site support isoprenoid chain elongation

According to the current model of chain elongation by *cis*-prenyltransferases, the pyrophosphate headgroups are bound at the superficial polar region, while the carbon chains point towards the deep hydrophobic tunnel. During the catalytic cycle, the pyrophosphate group of the initiatory substrate at the S_1_ site (FPP, C_15_) is hydrolyzed, followed by condensation of the remaining carbons with the IPP (C_5_) from the S_2_ site, yielding a 20-carbon polymer. Then, the elongated product translocates to the S_1_ site, where the growing carbon chain permeates deeper into the hydrophobic tunnel of the active site. Finally, a new IPP molecule binds to the S_2_ site, and the cycle repeats until the active site can no longer accommodate the long-chain isoprenoid (Kharel et al., 2006; Ko et al., 2001).

The structure reveals that the hydrophobic tunnel of DHDDS is formed by 2 α-helices (α2, α3) and 4 β-strands (βA, βB, βE, βF), similar to other *cis*-prenyltransferases (Kharel et al., 2006) (Figure 4A,B). It was previously suggested that the opening between α2 and α3 may be larger in DHDDS compared to short- and medium-chain *cis*-prenyltransferases leading to a larger diameter enabling the accommodation of longer products (Kharel et al., 2006). Indeed, in DHDDS, the distance between α2 and α3, measured between the Cα atoms of W64 and L104, is 19.5 Å (Figure 4B). In contrast, the distance between the corresponding positions in UPPS (F56 and E96), which synthesizes a 55-carbon isoprenoid, is only 15.3 Å (Guo et al., 2005).

**Figure 4.**
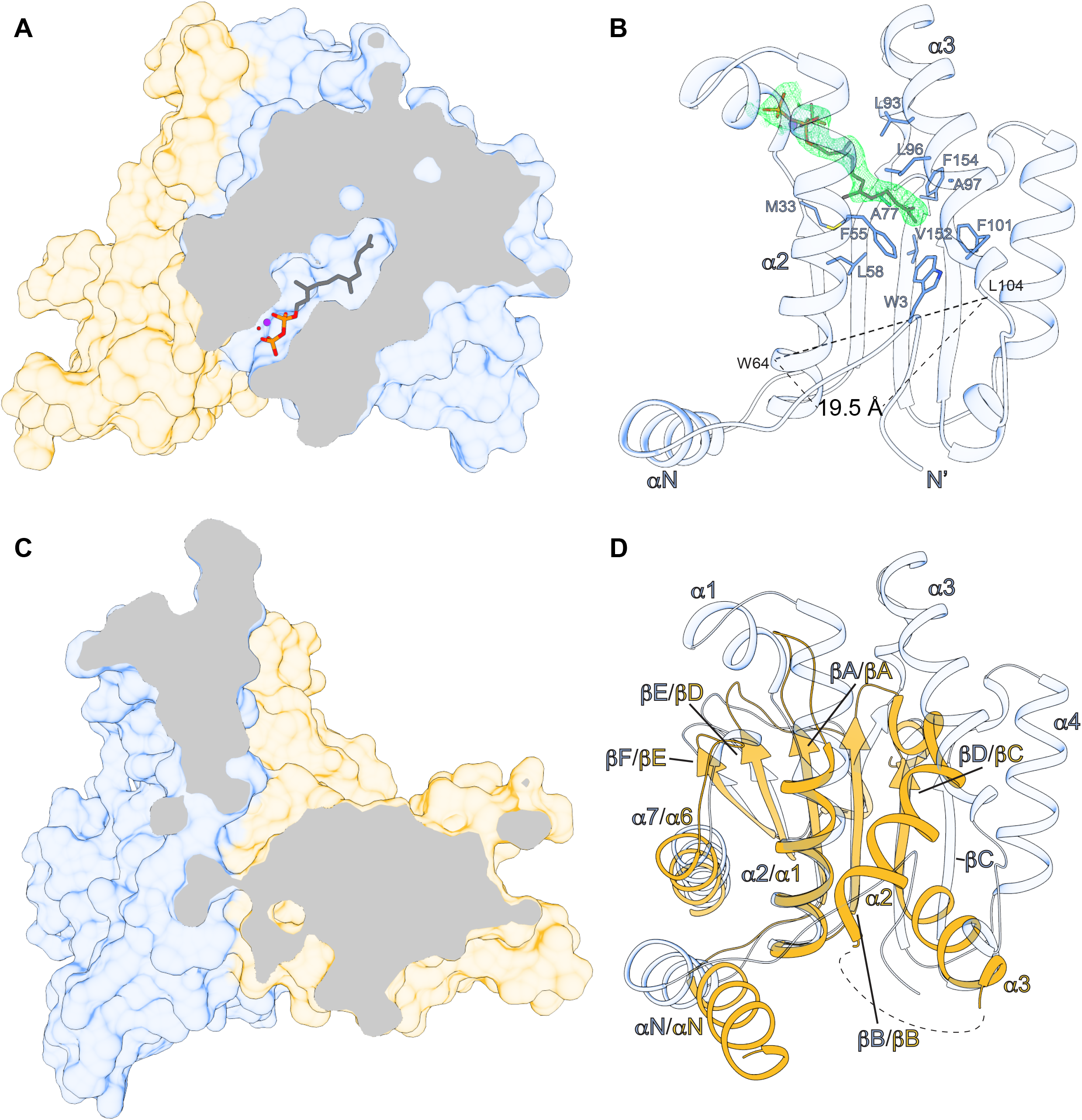
The hydrophobic active-site tunnel. **(A)** Clipped surface representation of the heterodimer showing the FPP binding site in DHDDS. Mg^2+^ and water molecules are shown as purple and red spheres, respectively. FPP is shown as sticks. **(B)** Coordination of the FPP tail within the active site. The 2F_o_-F_c_ electron density map of FPP, contoured at σ = 1, is presented as a green mesh. FPP interacting residues are shown as sticks. **(C)** Clipped surface representation of the heterodimer showing the lack of solvent accessible cavity in NgBR. **(D)** Cartoon representation of a superposition between the *cis*-prenyltransferase homology domains of DHDDS and NgBR. A collapse of the hydrophobic tunnel in NgBR is revealed. The secondary structure elements are indicated.

Intriguingly, the structure shows that the unique NTD of DHDDS can snake into the binding site, interacting with the bound FPP molecule (Figure 4B). Moreover, the N-terminal W3 interacts with F55, F101 and V152, thereby occluding the outlet of the hydrophobic tunnel of the active site. This results in a surprisingly small active site volume of 318 Å^3^. Indeed, compared with the 371 Å^3^ in active site of the medium-chain UPPS (Tian et al., 2018), the sh*cis*-PT site is seemingly inadequate for accommodating long-chain products. However, the high B-factors of the NTD (Figure S4) suggest that it is mobile, possibly assuming different orientations relative to the active site under physiological conditions. For example, the NTD may shield the hydrophobic tunnel in the apo or FPP-bound states, while being expelled from the tunnel upon chain elongation.

NgBR has been long known to lack catalytic activity of its own (Harrison et al., 2011). Nevertheless, the structural basis for this observation remained obscure. The structure clearly reveals that, in sharp contrast with DHDDS, sNgBR does not contain substrates binding sites (Figure 4C). Indeed, while the heterodimerization interface is structurally conserved (Figure 2A), the NgBR region corresponding to the active site in *cis*-prenyltransferases displays a significantly different structural arrangement (Figure 4D). Specifically, strands βΑ, βΒ, βD and helices α1 and α6 are tightly packed via hydrophobic interactions, leaving this region without a detectable substrate binding cavity and completely devoid of water molecules. This arrangement provides a structural explanation for the absence of NgBR catalytic activity, due to its incapacity to bind FPP and IPP. Thus, the only active site of the complex is situated within the *cis*-prenyltransferase homology domain of DHDDS (Figure 2).

### Active site-resident disease-associated mutations hinder the catalytic activity of h*cis*-PT

h*cis*-PT is a key player in dolichol synthesis, an essential moiety for the protein N-glycosylation process (Schwarz and Aebi, 2011). Underscoring its central functional importance, mutations in both complex subunits were recently associated with diverse human diseases, ranging from isolated blindness to fatal glycosylation disorders (Guo et al., 2018; Hamdan et al., 2017; Park et al., 2014; Sabry et al., 2016; Zelinger et al., 2011; Züchner et al., 2011). Mapping these mutations onto the structure reveals their intriguing clustering around the pyrophosphate binding regions of the S_1_ and S_2_ sites (Figure 5). Interestingly, with the exception of K42E, which leads to isolated retinitis pigmentosa, the rest of the mutations involve residues directly coordinating the pyrophosphate groups.

**Figure 5.**
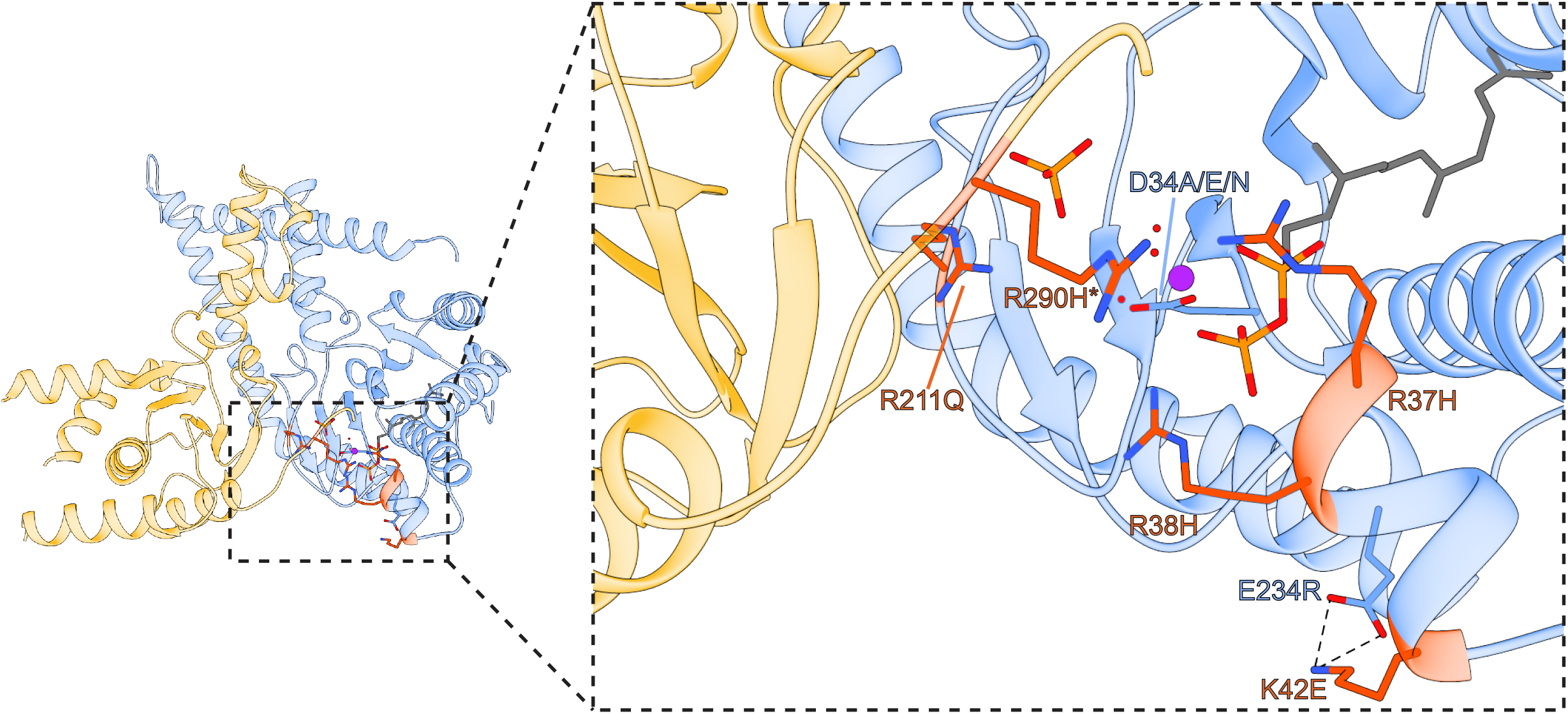
Structural basis for h*cis*-PT related diseases. Mapping of disease-associated mutations onto the sh*cis*-PT heterodimer. Disease-associated positions are presented as sticks and colored orange.

The S_1_ site mutations R37H and R38H, as well as the S_2_ site mutation R211Q, were previously associated with developmental encephalopathic epilepsies, highlighting the phenotypic convergence of mutations at the superficial polar region of the active site. Their direct involvement in substrate binding (Figure 3A, B) suggests that these three mutants may result in reduced catalytic activity. In addition to the DHDDS binding site mutants, a mutation of the conserved RxG motif of NgBR, R290H*, was shown to result in a congenital glycosylation disorder. Localized between the S_1_ and S_2_ sites (Figure 5), this residue does not form a direct interaction with either the phosphate molecule or the FPP. However, inspection of other *cis*-prenyltransferases structures, obtained in the absence of Mg^2+^, reveals that this conserved arginine can spatially and electrostatically replace the Mg^2+^ ion, transversely interacting with both the position corresponding to D34 and the pyrophosphate groups at S_1_ and S_2_ (Figure S5). Thus, in line with the high concordance between the spatial positioning and chemical properties of R290*, a histidine residue at this position is expected to result in functional perturbation (Grabińska et al., 2017; Park et al., 2014).

Although K42 does not interact with the substrate or binding site residues (Figure 5), it results in activity reduction, similar to R290H* (Grabińska et al., 2017). Careful examination of K42 surroundings revealed that it forms a salt bridge with E234 (Figure 5). Thus, we hypothesize that the K42E mutation results in repulsion between the positions, leading to a compensatory interaction of the mutant glutamate with adjacent positively charged active site residues such as R38 (Figure 5). These newly formed aberrant polar networks may in turn hinder the interaction of active site residues with the pyrophosphate groups, thereby reducing the catalytic activity. Together, disease-associated mutations are clustered in the proximity of the active site polar region, leading to direct interference with substrate binding and catalysis.

## Discussion

Here, we provide the first structure of a heteromeric *cis*-prenyltransferase. Our biochemical and structural analyses of h*cis*-PT reveal a novel heterotetrameric assembly, formed via a dimer-of-heterodimers mechanism, mainly through homotypic interface formation between the newly identified CTD in DHDDS (Figure 2). Furthermore, the structure elucidates how the architecture of NgBR precludes its endogenous catalytic activity while enabling allosteric intersubunit communication by transverse interactions of the conserved C-terminal RxG motif with the active site of DHDDS (Figures 3, 4). In addition, as the first long-chain *cis*-prenyltransferase structure, analysis of h*cis*-PT highlights the molecular determinants supporting formation of long-chain isoprenoid (Figure 4). Finally, the structure sheds light onto the mechanisms by which disease-associated mutations perturb catalytic activity (Figure 5).

To date, dimerization is considered as the common tertiary organization of *cis*-prenyltransferases (Fujihashi et al., 2001; Ma et al., 2019; Takahashi and Koyama, 2006). Additionally, the structurally characterized family members were homodimeric, encompassing an active site within each subunit and thus exhibiting an overall functional symmetry. In contrast, we show here that h*cis*-PT exhibits a novel heterotetrameric organization, achieved via a dimer-of-heterodimers assembly mediated by homotypic interactions of the CTD of DHDDS. Importantly, intact h*cis*-PT, expressed and purified from Expi293F cells, was previously subjected to size exclusion analysis. Although its mass was never directly assessed, it shares a markedly similar elution profile with sh*cis*-PT (Grabińska et al., 2017). This tetrameric assembly mode may contribute to the previously suggested mutual stabilizing effect resulting from NgBR and DHDDS co-expression in cells (Harrison et al., 2011).

The organization of the unique CTD of DHDDS, underlying h*cis*-PT tetramerization, provides a mechanistic explanation for the poor activity exhibited by DHDDS homodimers (Lisnyansky Bar-El et al., 2019). The helix-turn-helix motif, following α7 (Figure 2), is incompatible with formation of the extensive interactions network observed within each h*cis*-PT heterodimer active site (Figure 3). In contrast, the assembly of DHDDS with NgBR allows the complementation of the active site by the transverse interactions with the C-terminal tail of NgBR (Figure 3). Importantly, such transverse interactions are observed also in homodimeric *cis*-prenyltransferases (Figure S3), lacking a C-terminal helix-turn-helix motif, supporting the functional importance of these interactions in coupling active site organization with enhanced catalytic activity. Consistent with this notion, mutations in the C-terminal tail of NgBR were previously shown to result in drastically reduced catalytic activity (Grabińska et al., 2017; Park et al., 2014). Now, our structure offers a mechanistic understanding of the effects imposed by these mutations. Indeed, the G292A* mutation, previously introduced to explore the functional role of the conserved RxG motif, increased the K_m_ for IPP by ~6-fold while reducing the turnover rate by ~12 fold (Grabińska et al., 2017). As shown here, G292* is directly involved in IPP binding (Figure 3) and measuring the φ/ψ angles that this position can only be occupied by glycine (Ho and Brasseur, 2005). In addition, we show that K293* interacts with the catalytic residues R37 and R85 via its backbone carboxylate (Figure 3C). In accordance with the pivotal role of this interaction network, either deletion of K293 or the addition of a terminal alanine resulted in diminished catalytic activity of the intact complex (Grabińska et al., 2017). Together, the heterodimerization architecture observed here supports the notion that although DHDDS is considered as the catalytically active subunit, both subunits are necessary for efficient dolichol synthesis.

Our structure hints towards the mechanisms allowing long-chain isoprenoid synthesis by h*cis*-PT. While the deep hydrophobic tunnel, engulfing the elongating product, is walled by α2, α3, βA, βB, βE, βF (Figure 4), previous studies pinpointed the length of α3 as a key contributor to chain length determination (Kharel et al., 2006). Indeed, previous studies of UPPS showed that an insertion of 3 residues, corresponding to ^107^EKE^109^ in human DHDDS, led to an increase in product length from C_55_ to C_70_. The product length was further increased to C_75_ if a 5 residues insertion, mimicking the yeast ortholog Srt1, was introduced, establishing the correlation between the length of α3 and the product (Kharel et al., 2006). In full agreement with these observations, the structure of h*cis*-PT reveals that the ^107^EKE^109^ results in a kink in α3, leading to a ~4Å increase in the hydrophobic tunnel diameter (Figure 4B) compared with UPPS. In addition to the active site diameter, the composition of its terminal region also plays a key role in determination of product length. In UPPS, L137, localized to the N-terminus of βD, was shown to be vital for determining chain length, with the L137A mutant increasing product length from C_55_ to C_75_ (Ko et al., 2001). Our structure reveals that the corresponding position in DHDDS is replaced by C148, a hydrophilic and less bulky residue. Thus, C148 cannot occlude the hydrophobic tunnel outlet as efficiently as L137, similar to the L137A mutant. Together, the increased length of α3 and the composition of the hydrophobic tunnel outlet jointly contribute to the longer chain lengths produced by h*cis*-PT.

By forming h*cis*-PT, DHDDS and NgBR were shown to play a crucial role in cellular dolichol synthesis mechanism (Endo et al., 2003; Harrison et al., 2011). Mutations found in both subunits lead to a wide variety of devastating congenital syndromes (Hamdan et al., 2017; Park et al., 2014; Sabry et al., 2016; Zelinger et al., 2011; Züchner et al., 2011). These mutations cluster around the active site (Figure 5), which is mainly by DHDDS, with a key structural contribution from the C-terminus of NgBR (Figure 2,3). The structure provides a framework to better understand the mechanisms of dolichol synthesis, its disruption in disease and may enable design of novel therapies for these devastating conditions.

## Acknowledgments

We thank the staff of I03 at the Diamond Light Source for assistance with diffraction experimentation. This work was performed in partial fulfillment of the requirements for a Ph.D. degree of M.L-B., Sackler Faculty of medicine, Tel Aviv University, Israel. This work was supported by the Israel Science Foundation (grant number 1721/16) (Y.H.), the Israel Cancer Research Foundation grants 01214 (Y.H.) and 19202 (M.G.) and from the German☐Israeli Foundation for Scientific Research and Development grant number I☐2425☐418.13/2016 (Y.H.). Support also came from the I☐CORE Program of the Planning and Budgeting Committee and The Israel Science Foundation grant 1775/12 (Y.H.) and the Claire and Amedee Maratier Institute for the Study of Blindness and Visual Disorders, Sackler Faculty of Medicine, Tel-Aviv University (Y.H. and M.G.). Access to MS installation was funded by the EU Horizon 2020 grant EU_FT–ICR_MS project number 731077 and by CIISB LM2018127. P.M. and P.V. support from MEYS CZ funds CZ.1.05/1.1.00/02.0109 is gratefully acknowledged.

## Author Contributions

Conceptualization, Y.H. and M.G.; Methodology, Y.H. and M.G.; Investigation, M.L-B., P.V., P.M., Y.H. and M.G.; Formal Analysis, M.L-B., P.M., Y.H. and M.G.; Writing – Original Draft, Y.H. and M.G.; Writing – Review & Editing. M.L-B., P.M., Y.H. and M.G.; Supervision, Y.H. and M.G.; Funding Acquisition, Y.H. and M.G.

## Materials and methods

### Cloning

Full-length human DHDDS (residues 1-333, UniProt Q86SQ9) was cloned into pET-32b plasmid and sNgBR (residues 73*-293*, UniProt Q96E22) or sNgBRΔ167*-175* were cloned into pETM-11 plasmid as thioredoxin (TRX) fusion proteins, as previously described (Giladi et al., 2017). The constructs include a 6xHis-tag (DHDDS) or Strep-tag (sNgBR/sNgBRΔ167*-175*) to facilitate protein purification and a TEV-protease (Tobacco Etch Virus) cleavage site to remove the affinity tags and TRX fusion. Mutations were introduced using the QuickChange method and verified by sequencing.

### Protein expression and purification

*E. coli* T7 express competent cells were co-transformed with DHDDS and sNgBR (sh*cis*-PT) or sNgBRΔ167*-175* (x*cis*-PT), grown in Terrific Broth medium at 37°C until reaching OD_600nm_ = 0.6 and induced at 16°C by adding 0.5 mM isopropyl β-D-1-thiogalactopyranoside (IPTG). Proteins were expressed at 16°C for 16-20 h, harvested by centrifugation (~5,700xg for 15 min), and then resuspended in a buffer with 1 μg/ml DNase I and a protease inhibitor mixture. Resuspended cells were homogenized and disrupted in a microfluidizer. Soluble proteins were recovered by centrifugation at ~ 40,000xg for 45 min at 4°C. Overexpressed proteins were purified on a HisTrap HP column, followed by purification on a Strep-Tactin column and TEV protease cleavage of the purification tags and TRX fusions. The reaction mixture was concentrated and loaded onto a Superdex-200 preparative size-exclusion column pre-equilibrated with 20 mM HEPES, pH 7.5, 150 mM NaCl, 1 mM TCEP. Purified proteins were flash-frozen in liquid nitrogen and stored at −80° C until use. Protein purity was >95%, as judged by SDS-PAGE.

### Cross-linking

8☐μM of sh*cis*-PT were cross-linked by incubation with 0.005% glutaraldehyde at room temperature for 15 min. Reactions were quenched by the addition of sodium dodecyl sulfate and β-mercaptoethanol containing sample buffer, followed by 10☐min incubation at room temperature. Cross-linked products were analyzed by SDS-PAGE.

### ESI-MS

sh*cis*-PT (20 μM) was transferred into 200 mM ammonium acetate pH 7.5 by Zeba Spin columns (0.5 mL, 7-kDa cut off) and adjusted to 10 μM concentration. Sample was loaded into a home-made quartz-glass ESI tip which was mounted onto a custom built nESI source interfaced to Waters Synapt G2Si. Analyses at different activation settings were performed. Low activation (trap collision energy 10V) was used to obtain native-like conditions while high trap collisional energies (up to 110V) were employed to strip the adducts and obtain more accurate mass. Key instrument parameters were: sampling cone voltage 40V, source offset 20V, trap gas 4ml/min and source temperature 20 °C, ESI tip voltage 1.8kV. Data were analyzed in MassLynx 4.1.

### Enzyme kinetics

The activity of purified DHDDS was assayed as previously described (Edri et al., 2017; Giladi et al., 2017; Lisnyansky Bar-El et al., 2019). Briefly, 0.01-0.1 μM of purified proteins were mixed with FPP and [^14^C]-IPP to initiate the reaction in buffer composed of 25 mM Tris-HCl, pH 7.5, 150 mM NaCl, 10 mM β-mercaptoethanol, 0.02% Triton-X100, 0.5 mM MgCl_2_ at 30°C. 15 mM EDTA (final concentration) were added to quench the reaction and 500 μL of water-saturated 1-butanol was added to extract the reaction products by thorough vortexing. Initial rates were measured by quenching the reaction at 10% or lower substrate consumption. The K_m_ value of FPP was determined by varying [FPP] while holding [IPP] constant at 100 μM, and the K_m_ value of IPP was determined by varying [IPP] while holding [FPP] constant at 10 μM. The product, encompassing ^14^C, was quantitated using a scintillation counter. Kinetic constants were obtained by fitting the data to the Michaelis-Menten equation using Origin 7.0 (OriginLab, USA).

### Crystallization and structure determination

Initial crystallization screens were performed using ~15 mg/mL purified x*cis*-PT in the presence of 0.5 mM MgCl_2_ and 760 μM FPP at 19°C using the sitting drop vapor diffusion method. Crystals were obtained in 0.1 M NaCl, 0.1 M NaP pH 7.0, 33% w/v PEG 300. Data were collected at 100°K at the Diamond Light Source (DLS; Oxfordshire, United Kingdom). Integration, scaling and merging of the diffraction data were done with the XDS program (Kabsch, 2010). The structure was solved by automated molecular replacement and initial model building using the programs MrBump (Keegan and Winn, 2007) and CCP4 (Winn et al., 2011) (Table 1). Iterative model building and refinement were carried out in PHENIX (Adams et al., 2010) with manual adjustments using COOT (Emsley and Cowtan, 2004). Structural illustrations were prepared with UCSF Chimera (https://www.cgl.ucsf.edu/chimera). Atomic coordinates and structure factors for the structure of x*cis*-PT in complex with Mg^2+^ and FPP have been deposited in the Protein Data Bank with accession number 6Z1N.

**Table 1.** Crystallographic statistics.

| **Data collection** |  |
| --- | --- |
| Beamline | DLS I03 |
| Wavelength (Å) | 0.976 |
| Space group | R32:H |
| Cell dimensions |  |
| $a, b, c$ (Å) | 184.1, 184.1, 112.6 |
| $\alpha, \beta, \gamma$ (°) | 90, 90, 120 |
| Resolution (Å) | 2.30 (2.44-2.30) |
| R<sub>meas</sub> (%) | 17.0 (197.8) |
| CC<sub>1/2</sub> (%) | 99.9 (67.7) |
| Completeness (%) | 100.0 (100.0) |
| Mean I/σ(I) | 16.3 (1.6) |
| Multiplicity | 21.1 (20.7) |
| **Refinement statistics** |  |
| No. reflections (work/free) | 30844/1633 |
| Resolution range | 45.99-2.30 |
| R<sub>work</sub>/R<sub>free</sub> | 0.1895/0.2209 |
| No. atoms |  |
| Macromolecules | 4,079 |
| Ligands | 30 |
| Solvent | 122 |
| Average *B*-factor |  |
| Macromolecules | 56.29 |
| Ligands | 57.25 |
| Solvent | 54.43 |
| RMSD (bond lengths) | 0.006 |
| RMSD (bond angles) | 1.03 |
| Ramachandran outliers (%) | 0 |

Values in parentheses are for highest-resolution shell

