## Supplemental information text for "Structure of the human heterotetrameric *cis*-prenyltransferase complex"

**Figure S1. Topology of sh*cis*-PT.** The secondary structures of DHDDS (A) and sNgBR (B) are indicated above the sequence.

**Figure S2. Native ESI analysis of sh*cis*-PT.** The calculated mass of the heterotetramer using high activation conditions is 128.7 ± 0.02 kDa.

**Figure S3. IPP coordination at the S_2_ site of homodimeric *cis*-prenyltransferase.** The IPP pyrophosphate binding region at the S_2_ site of the homodimeric *mycobacterium tuberculosis* decaprenyl diphosphate synthase (PDB 6IME). One monomer is colored green and the other pink. IPP and its coordinating residues are shown as sticks. The organization of the binding site and the interaction network with IPP is identical to that observed for the phosphate molecule in the DHDDS active site.

**Figure 4. B-factors plots.** Mean residue B-factors of DHDDS (left) and NgBR (right). The black line represents the average macromolecule B-factor.

**Figure 5. Transverse interactions of R290^*^ with the active site.** Cartoon representation of the pyrophosphate binding region of sh*cis*-PT (left) and the homodimeric *mycobacterium tuberculosis* farnesyl diphosphate synthase (PDB 2VG2) (right), colored by chain. Note the side-chain movement of R292**^*^**, corresponding to R290**^*^** of NgBR, to replace the Mg^2+^ ion (right).
