## Supplementary figures and images for "Structure of the human heterotetrameric *cis*-prenyltransferase complex"

### Figure S1

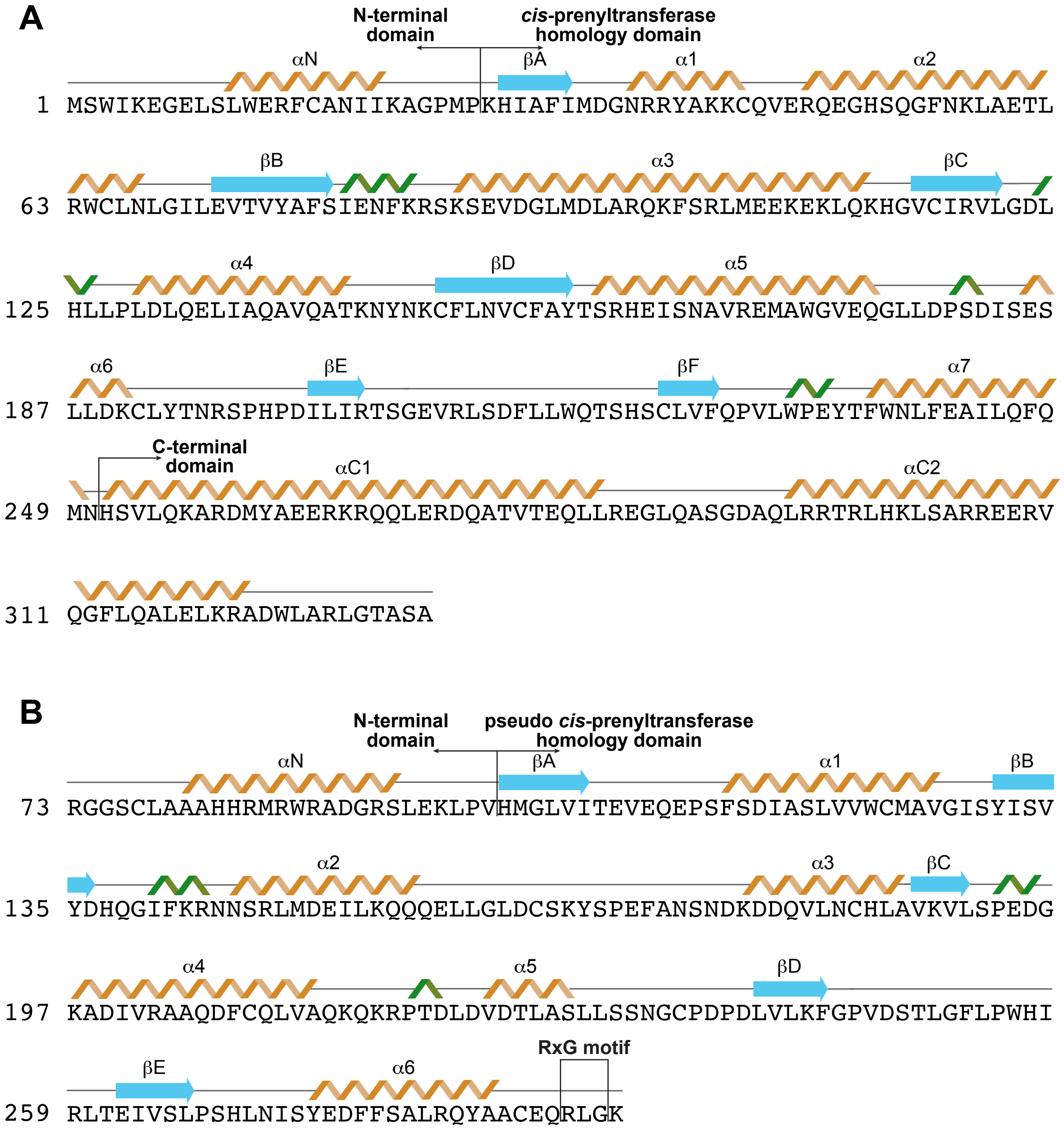

### Figure S2

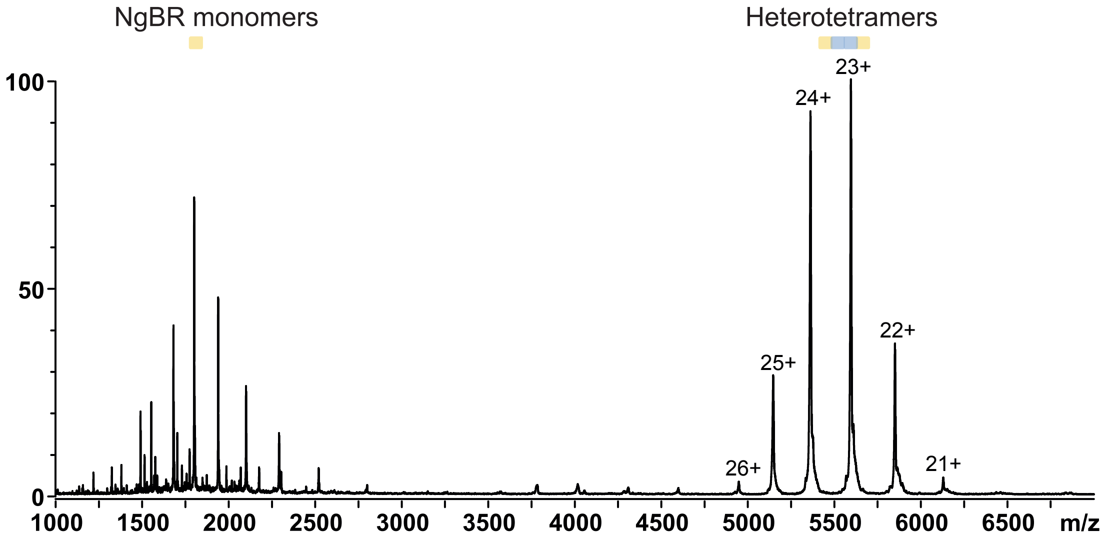

### Figure S3

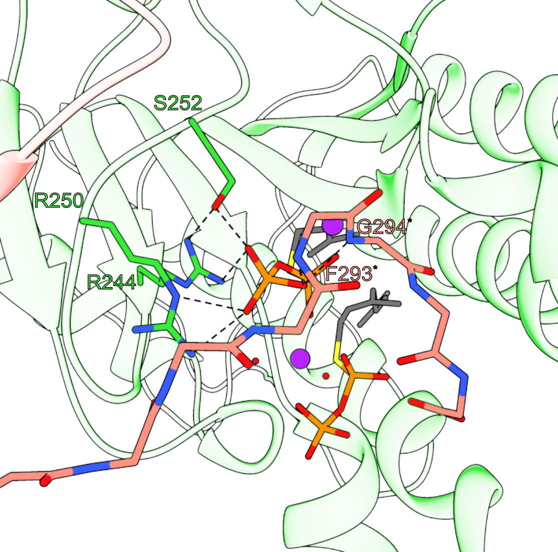

### Figure S4

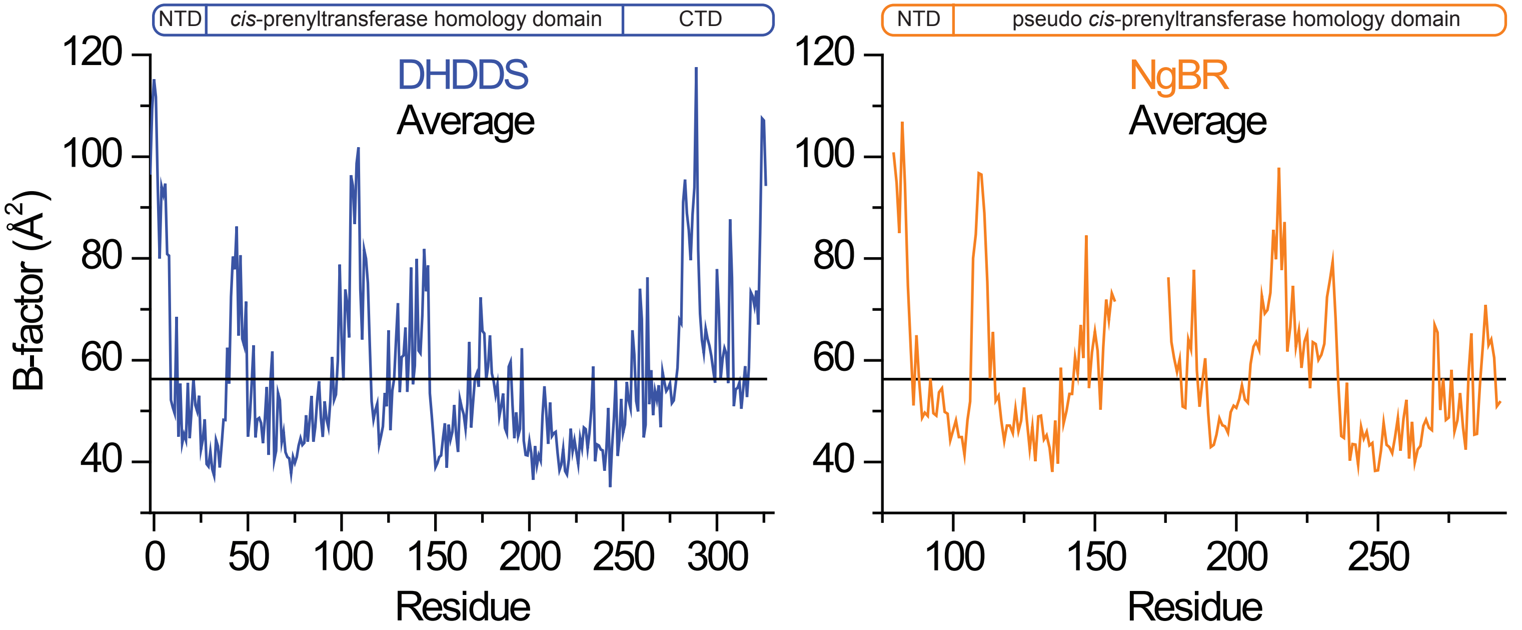
